# The Research Landscape of Dynamic Functional Connectivity in Parkinson’s Disease: A Scoping Review and An Interactive Tool

**DOI:** 10.64898/2025.12.08.692999

**Authors:** Daniel Kristanto, Daniela Rodriguez De Castro, Amirhussein Abdolalizadeh

## Abstract

Dynamic functional connectivity (dFC) analysis of functional Magnetic Resonance Imaging (fMRI) data has emerged as a powerful framework for characterizing the time-varying network disintegration and topological transitions underlying Parkinson’s disease (PD). However, the rapid expansion of this field introduces methodological heterogeneity, creating a fragmented landscape that complicates knowledge synthesis and informed decision-making for future study designs. To address this, we performed a scoping review to map the conceptual and methodological diversity of dFC research in PD. Beyond a traditional static synthesis, we developed DynaPD, an open-source interactive application that allows researchers to dynamically explore study designs, analytical pipelines, and reported findings. Our synthesis of 37 eligible studies reveals divergence in preprocessing strategies but a convergence on core analytical pipelines, notably sliding-window techniques coupled with k-means clustering. Empirically, studies converge on the importance of neural flexibility assessed via dwell times and topological transitions between strongly and sparsely connected states. However, specific network interpretations remain variable. Common limitations included cross-sectional designs, limited spatial resolution for subcortical networks, and short scan duration. By providing a scoping evidence synthesis and a living interactive tool, this work functions as a decision-support resource to consolidate the literature and guide the methodological design of future studies.

## 1. Introduction

Parkinson’s disease (PD) is a progressive neurodegenerative disorder increasingly understood as a profound disruption of large-scale brain networks ^1,2^. While characterized by complex motor and cognitive decline, these symptoms are not merely attributable to isolated regional dysfunction but rather reflect widespread network disintegration ^3^. In network neuroscience, topology refers to the architectural arrangement of brain regions and their connections. Topological shifts denote large-scale reconfigurations in this architecture, for example, the transition from globally integrated networks to hsegregated, isolated sub-networks. Mapping these shifts is therefore important for understanding the neural underpinnings of disease progression.

One of the most prominent research avenues is the use of functional Magnetic Resonance Imaging (fMRI) to characterize brain profiles in PD. Static Functional Connectivity (sFC), which measures time-averaged co-activation between brain regions, became a widely adapted measure. For example, sFC studies have associated cognitive decline in PD with an overall increase in local Functional Connectivity (FC) and a reduction in long-range FC, indicating a loss of optimal network topology^4^. Similarly, weaker intra-network FC within the Fronto-Parietal Network (FPN) has been linked to lower cognitive scores in PD ^5^. A meta-analysis of α-Synucleinopathies also characterized PD by decreased FC in the Default Mode (DMN) and Ventral Attention Networks (VAN), which are important for cognition and attention^6^.

Despite these insights, sFC assumes that functional relationships are stable, representing time-averaged connectivity over several minutes. This static approach cannot capture the temporal flexibility of the brain, which is key for understanding the dynamic interplay between motor and cognitive systems. For instance, the co-occurrence of motor and cognitive symptoms may be explained by a neural compensation framework, where the brain actively recruits additional cortical resources, particularly from cognition-related networks, to offset dysfunction in motor-related networks ^7^. Capturing such time-varying processes requires a dynamic measure.

In contrast, dynamic Functional Connectivity (dFC) analysis captures connectivity patterns as they evolve over time, offering a window into neural flexibility and topological transitions^8,9^. Given this potential, the implementation of dFC is growing in PD research. For instance, Fiorenzato et al. (2019) used a sliding-window approach to characterize PD. They found that dFC in PD patients was represented by a more frequent, weakly-connected brain state and a less frequent, strongly-connected brain state. Moreover, a reduced number of transitions between states was associated with worse cognitive decline. The dFC framework thus holds significant promise for elucidating the neural underpinnings of parallel cognitive and motor decline in PD.

However, modeling time-varying brain networks introduces significant computational challenges, leading to considerable heterogeneity in study designs and analytical pipelines. This variability complicates the synthesis of current knowledge and may hinder replicability^11^. Therefore, it is necessary to map this heterogeneity to clarify the current research landscape and inform future studies. To address this, the present scoping review aims to systematically map and visualize the methodological and conceptual landscape of dFC-fMRI research in PD. The most related review was published in 2019 about dFC in neurodegenerative disorders ^12^. In this review, PD was one of the topics; however, the content was limited and focused more on the findings, without discussing the vast variability of study designs and computational frameworks addressed in the current study. Moreover, the present review also offers an interactive, web-based application to explore the findings. This tool serves as a complementary component of our review, enabling users to dynamically explore the extracted information, filter for studies matching specific criteria (e.g., methods or clinical focus), and investigate the high degree of variability in analytic decision-making. Taken together, this scoping review and its accompanying interactive tool aim to map the research landscape of this field, making dFC research in PD more coherent and promoting an informed, integrative approach for future study designs.

## 2. Methods

### 2.1. Specific aim of the review

This scoping review aims to map the methodological and conceptual landscape of dFC fMRI research in PD. As this field rapidly expands, it has produced a diverse body of literature with substantial variability in computational approaches and topological findings. To address this, we moved beyond traditional static figures by introducing an interactive open-source application. Our specific objectives are:

1. To extract and synthesize key data points across studies, including sample characteristics, methodological pipelines, and reported network findings.
2. To interactively visualize the variability in analytical decision-making (e.g., in motion correction, dFC method, and state definition).
3. To interactively map the conceptual structure of the field, allowing researchers to explore the relationships between studies based on shared methodology, clinical focus, or findings.
4. To provide a dynamic tool that allows researchers to navigate this landscape, filter for studies matching specific criteria, and make more informed decisions for future analysis plans.

### 2.2. Search strategy

To ensure transparency in our study selection process, we adapted the flow diagram structure from the Preferred Reporting Items for Systematic reviews and Meta-Analyses extension for Scoping Reviews (PRISMA-ScR)^13^. The diagram illustrating our complete search and selection workflow is presented in Figure 1.

**Figure 1.**
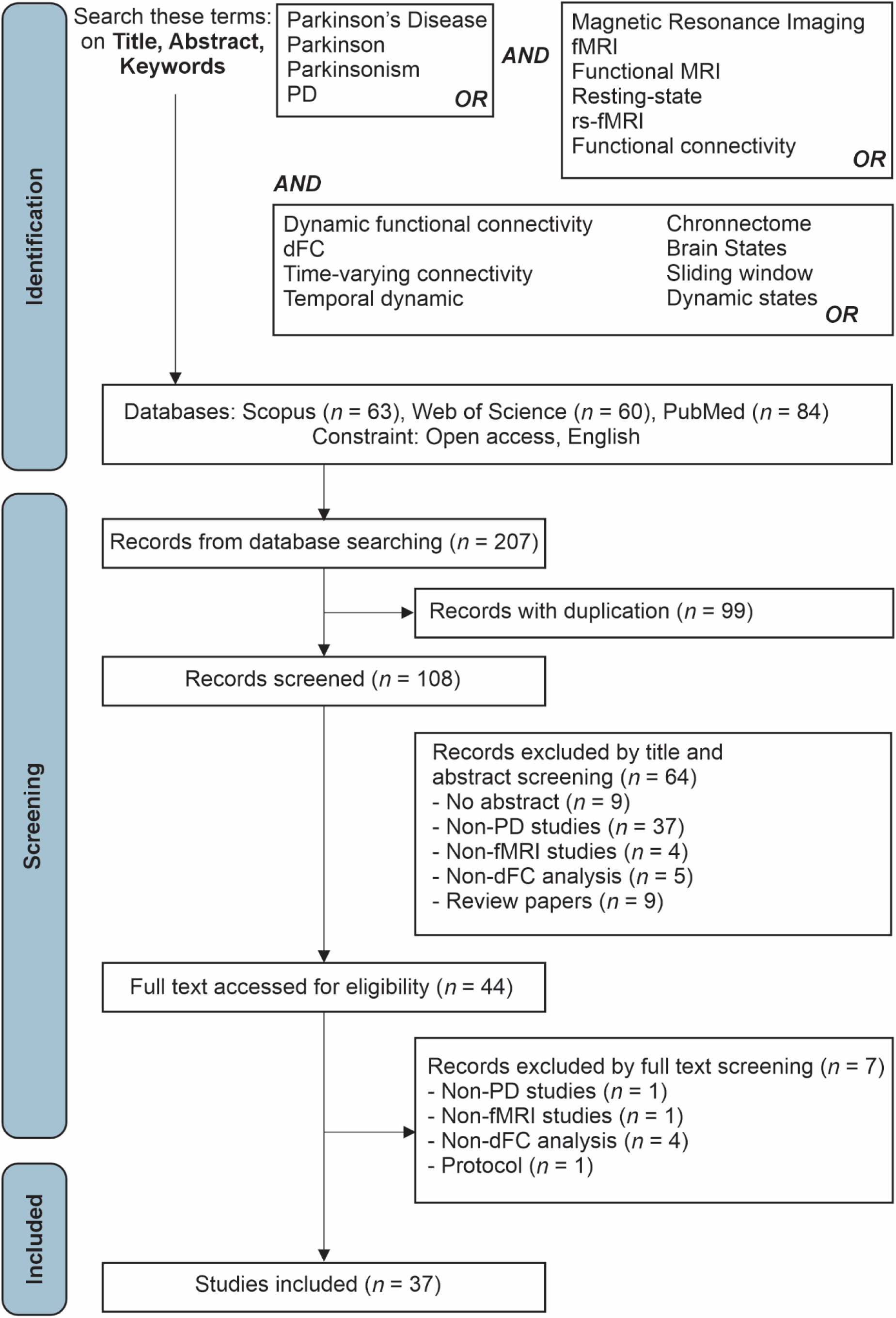
PRISMA diagram to report the identification, screening, and selection process of the studies included in this review.

As the scope of this review encompasses studies implementing dFC derived from fMRI in individuals with PD, our search query included terms related to PD, fMRI, and dynamic FC (see Figure 1 for the complete list of search terms). We limited the search to open-access articles published in the English language. This criterion was selected to align with the open-science framework of the accompanying interactive application, ensuring that all synthesized data and primary sources remain fully accessible to the community. The search was performed across three databases – Scopus, Web of Science, and PubMed – on June 6, 2025, yielding a total of 207 articles. Metadata for these articles were downloaded from each database in .bib format.

The .bib files were subsequently preprocessed using the R software for statistical computing, specifically employing the *revtools* package ^14^. First, we read and merged the .bib files across databases, transforming them into a single data frame using the `read_bibliography()` function. Next, we utilized the `find_duplicates()` function to identify duplicate items within the data frame, employing both DOI- and title-based matching approaches. We then removed the duplicates and retained unique articles using the `extract_unique_references()` function. This process resulted in 108 unique articles eligible for the screening phase. The scripts used for data preprocessing and dataset containing the extracted information are publicly available at [https://github.com/kristantodan12/dFC-PD].

### 2.3. Screening phase and eligibility criteria

The unique articles, organized in a data frame, were subjected to an initial screening phase based on their abstracts. This process was conducted using the `screen_abstract()` function from the revtools package. Specifically, we utilized the graphical user interface to manually review each abstract. A key advantage of this function is its capacity to allow users to annotate abstracts as ’included’ or ’excluded’ and to append specific notes to each entry. During this abstract screening phase, we excluded 64 articles from the initial pool of 108 unique studies. The most frequent rationale for exclusion (*N* = 37) was that the studies did not analyze a PD dataset. A detailed summary of the exclusion criteria is provided in Figure 1. Following this procedure, 44 studies proceeded to the subsequent eligibility screening phase.

To assess the eligibility of these studies, we first retrieved their full texts using Zotero (www.zotero.org), a reference management software. We compiled a list of Digital Object Identifiers (DOIs) for the candidate studies and imported this list into Zotero for automated full-text retrieval. Importantly, because our initial search strategy targeted open-access articles, Zotero successfully retrieved the full texts for all 44 studies. We then evaluated these studies against our inclusion criteria. This comprehensive assessment resulted in a final set of 37 articles included in this scoping review.

### 2.4. Data extraction procedure and quality assessment

As stated above, the primary aim of this scoping review is to map the conceptual and methodological landscape, as well as the network-based findings, of studies implementing fMRI-derived dFC in PD. To achieve this, we extracted a comprehensive range of information from the included studies, categorized into four main domains. The first category, *Publication Information* (e.g., authors, journal), was retrieved from the bibliographic metadata. Second, we extracted information regarding *Study Design*, including sample characteristics for both PD and Healthy Control (HC) groups, as well as the primary focus of the investigation. The third category, *Methodological Details*, encompassed the coding of fMRI preprocessing procedures (e.g., motion regression, temporal filtering) and dFC analysis pipelines (e.g., computational methods and parameters). Finally, we captured information related to *Findings*, specifically the number of identified states, the qualitative interpretation of these states, and the reported limitations. The complete dataset extracted from the 37 eligible studies is available in the Database.xlsx file at our GitHub repository [https://github.com/kristantodan12/dFC-PD], which also contains detailed descriptions for each extracted variable.

The extraction of information from the eligible studies was conducted by two independent coders (DK and DC). To ensure data quality, the information extracted by both coders was rigorously compared item-by-item (see Database.xlsx in the GitHub repository). Out of a total of 5,074 items coded (spanning 86 variables across 59 analyses from 37 studies), there were 74 discrepancies, resulting in an initial inter-coder agreement rate of 98.5%. To resolve these discrepancies, we performed a collaborative discussion to reach a consensus on the final coding. Following this harmonization, author DK re-verified the coded information for all articles to ensure consistency.

### 2.5. Interactive app for data visualization

A central aim of this review is also to enable the interactive exploration of our dataset, comprising the coded information from all eligible studies. To facilitate this, we developed an interactive visualization tool using the Shiny package ^15^ within the R statistical computing environment, similar to our previous study^16^. Designed to complement the static figures presented in this paper, this application allows users to dynamically navigate the landscape of fMRI-derived dFC research in PD, exploring dimensions of study design, sample characteristics, methodological details, and findings. Access to the application is available via two methods. First, users can clone the source code from our GitHub repository [https://github.com/kristantodan12/dFC-PD] and run the application locally by following the provided instructions. Alternatively, a live web-based version is available for immediate access. The active URL for this deployment is available in the GitHub repository. For the remainder of this paper, this application is referred to as DynaPD.

DynaPD has several tabs serving different features:

1. *Landing Page (Home):* Upon accessing DynaPD, users will find an interactive network graph representing the architecture of the app. Different nodes correspond to specific application tabs. Clicking a node provides a brief description of its contents and routes the user directly to that section.
2. *Full Dataset:* This section provides access to the underlying data structure and is divided into three subtabs. The *Dataset* subtab displays the PRISMA flow diagram, detailing the study selection process. The *Data Explorer* subtab features a highly interactive table of the complete dataset in long format. It includes column-filtering tools with dynamic range sliders for numeric variables and dropdown menus for categorical variables, allowing users to identify specific study subsets. Finally, the *Coded Information* subtab provides detailed descriptions of every extracted variable.
3. *Study Overview:* This tab offers a comprehensive summary of the included literature, covering publication metadata (year, journal), sample characteristics, and data sources. It incorporates dynamic filtering functionality, allowing users to refine the visualizations based on criteria such as publication year, PD sample size, study design, primary focus, and focus specification. A tabulated list of the filtered studies is provided below the visualizations to enable detailed exploration of individual papers.
4. *Method Explorer:* This section is divided into three subtabs: *MRI Acquisition and Preprocessing*, *dFC Analysis*, and *Clinical Focus and Brain Mapping*. The first subtab visualizes the variability in fMRI preprocessing pipelines, including motion regression strategies and exclusion criteria. The *dFC Analysis* subtab illustrates the landscape of methodological choices specific to dFC, such as brain mapping techniques, parcellation schemes, and clustering algorithms. Both subtabs are integrated with filtering functions similar to the Study Overview, but are further enhanced with range sliders for Repetition Time (TR), allowing for targeted methodological inquiries. The final subtab, *Clinical Focus and Brain Mapping*, enables users to identify which brain areas or networks were investigated in relation to specific clinical foci. Users can select a clinical focus from a dropdown menu, and the associated brain regions are visualized based on citation frequency.
5. *Finding Explorer:* This tab facilitates the exploration of variability in reported results and is organized into four subtabs. The *State Features* subtab visualizes the frequency of different dynamic features computed across studies. The *State Interpretation* subtab provides a detailed characterization of the interpretations and clinical implications of these dynamic states. To accurately capture nuanced clinical findings, it utilizes categorized bar charts to visualize the consensus (or lack thereof) regarding whether strongly-connected states are beneficial and sparsely-connected states are detrimental. The *Proposed Biomarkers* subtab presents a searchable table summarizing the specific dFC-based biomarkers proposed by each study. Finally, the *Limitations* subtab aggregates and visualizes the limitations most frequently cited across the reviewed literature.
6. *Network of Studies:* This tab allows users to explore the conceptual and methodological interconnectivity of the field. It generates network graphs showing how studies are linked based on various similarity criteria, including primary clinical focus, data source, investigated brain networks, dFC methods, or the specific state features computed.
7. *Contribute*: To ensure DynaPD remains a living resource that evolves with the field, this tab provides a structured pipeline for community contribution. Researchers can download a standardized CSV template, input the coded variables for newly published dFC-PD studies, and upload the file directly to the authors’ cloud database for verification and subsequent integration into the live application.

### 2.6. AI tool use statement

During the preparation of this work, the authors used Gemini Advanced (Google) to improve the readability, spelling, and grammar of the manuscript. After using this tool, the authors critically reviewed and edited the content and take full responsibility for the final content of the publication.

## 3. Results

The Results section is organized into four thematic domains. First, we present a comprehensive inventory of the information extracted from the included studies. The subsequent three sections map the landscape of study design, methodological approaches, and findings as well as study limitations. Importantly, while this section utilizes static figures to summarize key trends, we encourage readers to concurrently utilize the DynaPD application to explore granular data points and relationships not fully captured in the text. This interactive approach substitutes the need for a static summary table, allowing for dynamic filtering of all 37 included studies.

### 3.1. General overview and list of the coded information

In total, we identified 37 eligible studies, revealing a distinct upward trend in publication frequency in recent years. The most prominent publication venues included *Frontiers in Aging Neuroscience*, *Neurobiology of Disease*, and *NeuroImage: Clinical*. Across all studies, we aggregated data from a total of 1,941 participants with PD, with the majority of individual studies reporting sample sizes between 20 and 30 participants (see also the *Study Overview* tab in DynaPD).

Regarding the scope of extraction, we coded 85 distinct variables for each article, covering study metadata, experimental design, methodological specifications, and reported network findings. The complete list of these variables is available in the *Database.xlsx* file in our GitHub repository and accessible via the *Full Dataset* tab in the DynaPD app. It is important to emphasize that the high dimensionality of the extracted dataset is partly attributable to the lack of standardized reporting practices. For instance, demographic and clinical variables such as age and motor symptoms severity scores were reported inconsistently, ranging from mean and range (min/max) to mean and standard deviation, or median and interquartile range. Similarly, the granularity of computational fMRI preprocessing descriptions varied substantially across studies. Furthermore, regarding data accessibility, the majority of studies utilized private datasets. Among them, only two studies explicitly deposited their data in public repositories (OpenNeuro and Zenodo), 14 stated that data would be available upon request, one indicated future public availability, while the remaining studies did not specify any data availability statement (see also Figure 2).

**Figure 2.**
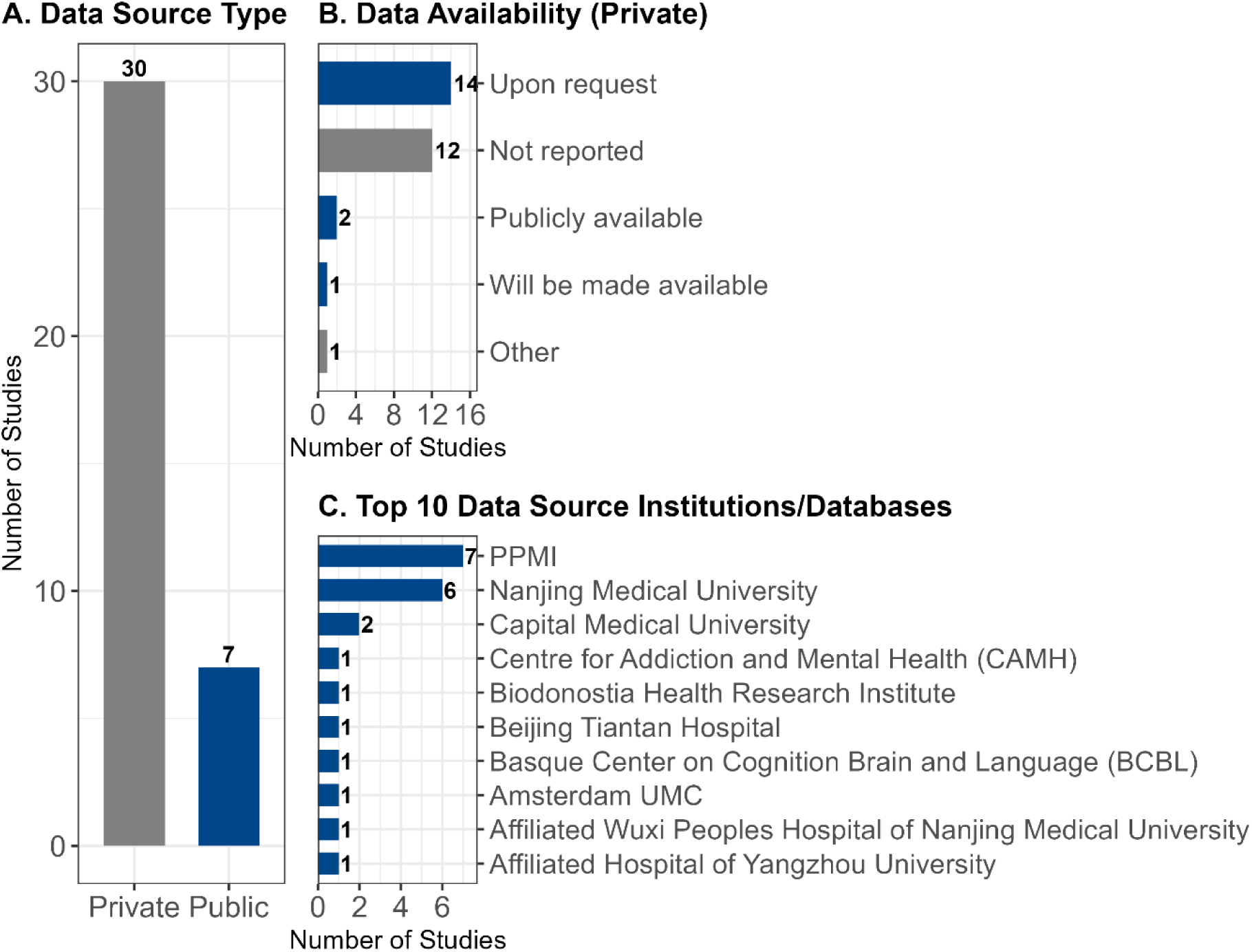
Data availability across studies. (A) The majority of studies utilized private datasets (*N*=30), with a smaller subset leveraging public repositories (*N*=7). (B) Among studies using private data, data availability policies varied: 14 studies stated data would be available upon request, while 12 studies did not specify any data sharing policy. Only one study indicated that private data would be made publicly available. (C) The PPMI (Parkinson’s Progression Markers Initiative) and Nanjing Medical University were the most frequently utilized data sources.

### 3.2. A cross-sectional and resting-state dominance with heterogeneous clinical targets

As illustrated by the Sankey diagram in Figure 3, the eligible studies are characterized by a predominance of cross-sectional designs targeting a heterogeneous array of clinical symptoms, including apathy, depression, freezing of gait, hallucinations, and sleep disorders. However, it is important to note that while these studies prioritized distinct non-motor foci, the assessment of motor severity (specifically MDS-UPDRS Part III) was a common feature across all investigations, serving as a standard baseline measure. Moreover, only 10 studies employed a longitudinal design; these focused primarily on disease progression and responses to diverse interventions, such as Deep Brain Stimulation (DBS) and dopaminergic medication (e.g., levodopa). This cross-sectional dominance fundamentally limits the ability to track topological network shifts over time. Furthermore, the resting-state paradigm was largely favored. Only three studies utilized task-based fMRI paradigms, specifically a modified Sternberg working memory task, an attention network test, and a right-hand squeeze-bulb motor modulation task. While resting-state fMRI provides a method for mapping baseline intrinsic network architecture, this methodological homogeneity may limit our understanding of state-dependent network flexibility. Parkinson’s disease is clinically characterized by deficits in initiating and switching between motor and cognitive states^3^. The lack of task-based dFC studies therefore represents a gap in the literature to directly observe how large-scale network topology dynamically reconfigures in response to external demands. Incorporating task-based paradigms into future dFC frameworks will be important for explicitly linking intrinsic network disintegration to external stimuli and cognitive impairments. To explore these trends further, readers are invited to filter studies by design and clinical focus within the *Full Dataset* tab of DynaPD. Additionally, this filtering functionality is integrated into most subtabs of the application, enabling a targeted exploration of the methodological and empirical landscape specific to a reader’s clinical interest or study design preference.

**Figure 3.**
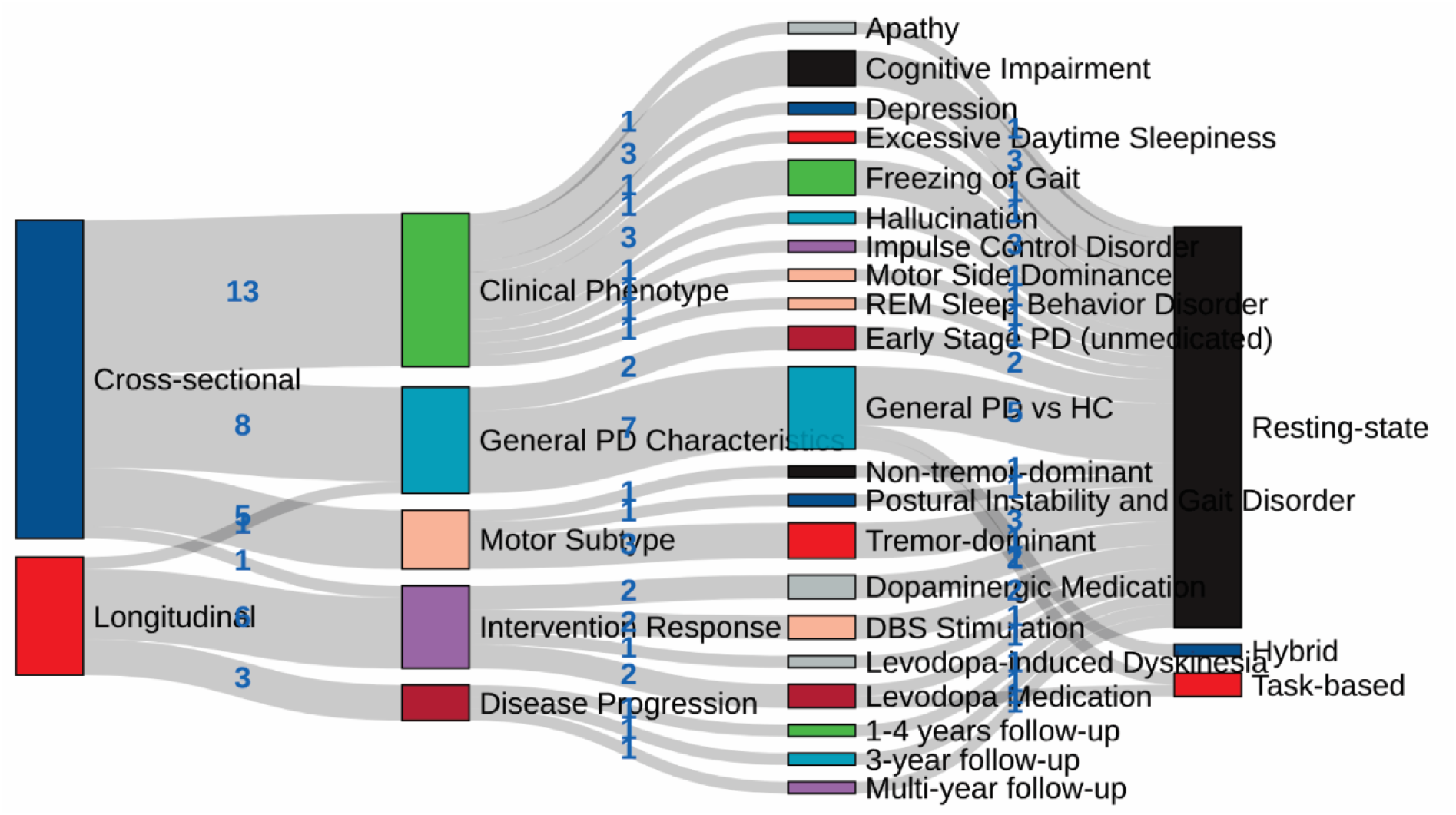
Landscape of study design, clinical focus, and MRI paradigms. The Sankey diagram visualizes the relationships between study designs, primary research foci, and fMRI paradigms. The majority of studies employed a cross-sectional design, which was applied to a broad and heterogeneous range of clinical targets, including general PD characteristics, motor symptoms, and specific non-motor symptoms (e.g., apathy, cognitive impairment). In contrast, longitudinal designs were less frequent and predominantly targeted disease progression and intervention response. Notably, regardless of the study design or clinical focus, the field demonstrates a reliance on the resting-state paradigm, with only a marginal number of studies utilizing task-based fMRI.

### 3.3. Methodological divergence in acquisition and denoising, followed by convergence on a common dFC analysis framework

We found significant variability in the methodological aspects of the reviewed studies. As depicted in Figure 4, the scanning protocols showed considerable heterogeneity, particularly regarding TR and scan length. While this variability may reflect resource constraints, it has important implications for computational analysis; referring to methodological benchmarks, datasets exceeding 10 minutes are recommended for robust dFC analysis, especially when using sliding-window approaches ^17^. Furthermore, we observed diverse strategies for fMRI data cleaning, with detrending being the most frequently applied step. Motion regression strategies also varied, with the 24-parameter model being the most popular choice, followed closely by 6-parameter and 12-parameter approaches.

**Figure 4.**
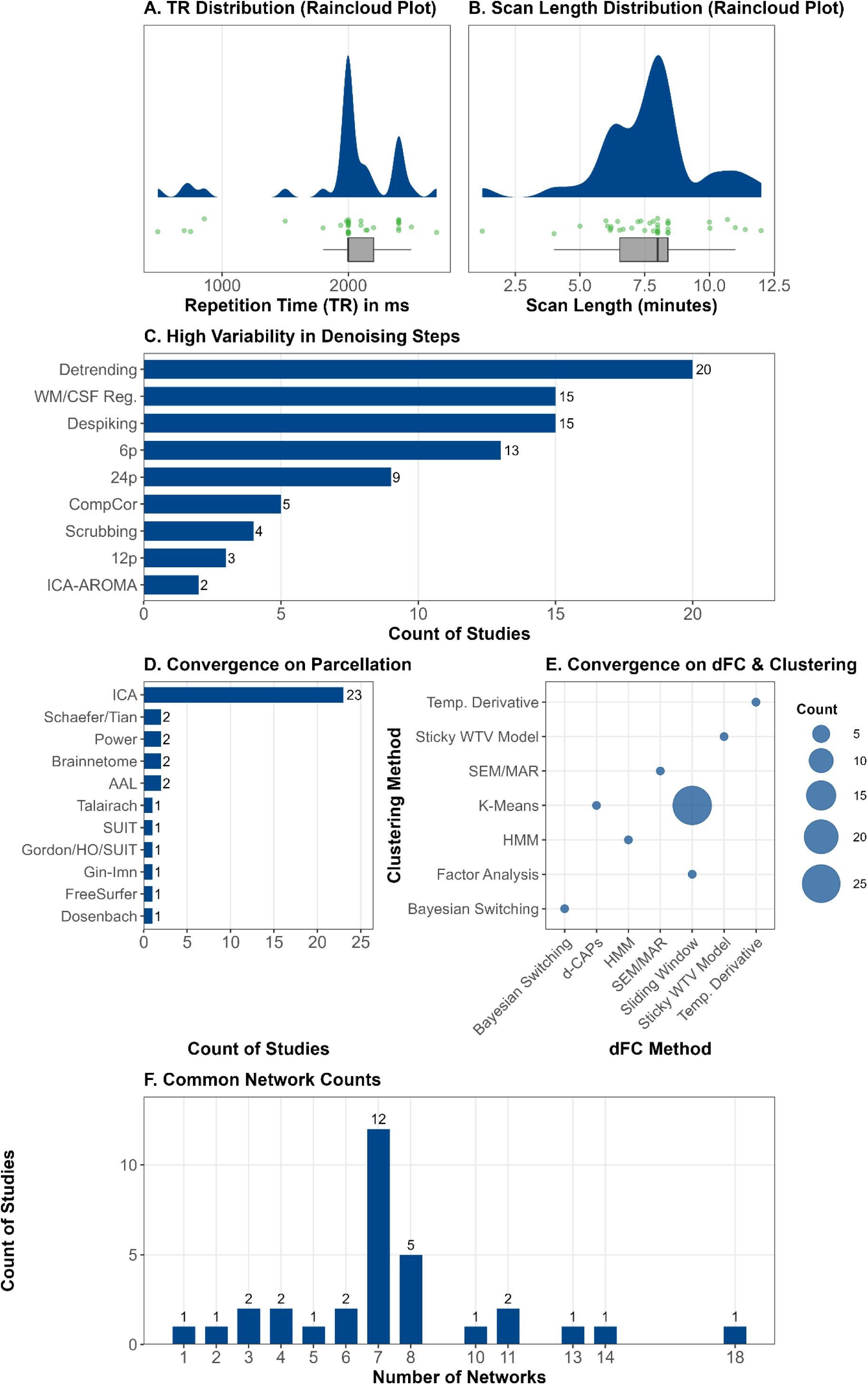
Methodological divergence, analytic convergence. (A-C) Divergence: Acquisition and preprocessing steps show high variability. TRs (A) and scan lengths (B) are broadly distributed, while preprocessing (C) reveals a diverse combination of motion regression parameters (24p, 6p, 12p) and other denoising steps. (D-F) Convergence: Analytic methods show strong consensus. ICA (D) is the most frequent parcellation choice, and the combination of sliding window and k-means (E) forms the dominant analytic core. Studies using network analysis also commonly converge on 7 networks (F).

In contrast to the divergence in preprocessing, post-processing steps demonstrated a convergence on specific methodological choices. To identify the regions or networks of interest, Independent Component Analysis (ICA), a data-driven technique, was the predominant approach. Similarly, the core dFC analysis pipeline showed strong consensus: the sliding-window technique was almost universally used for segmenting time-series, while k-means clustering served as the standard method for identifying topological brain states. Regarding network selection, most studies focused on the DMN and Sensorimotor (SMN) networks. Importantly, these methodological trends are fully explorable within DynaPD, where readers can dynamically adjust filters to investigate how methodological decisions vary across different studies.

### 3.4. Dynamic FC findings: consensus on state features, variable interpretations, and a common path forward

Figure 5 summarizes the key findings across the reviewed studies. Regarding state identification, the majority of studies reported a small number of distinct states, typically ranging from two to four. Importantly, regardless of the total number of states, a consistent dichotomy in connectivity patterns emerged: one characterized by weak, sparse inter-network connections (here is termed as sparsely connected states), and the other exhibiting strong, dense connections (here is termed as strongly connected states). Following state identification, studies typically computed specific dynamic features to investigate their associations with clinical or behavioral outcomes. The most commonly calculated metrics were: number of transitions between states; dwell time, representing the average duration of a state per visit; and fraction time, representing the total proportion of the scan duration spent in a specific state. Notably, a consensus emerged regarding the temporal dynamics of these patterns: strongly connected states were consistently found to be less frequent (or transient) compared to the predominant sparsely connected states.

We further synthesized the findings regarding the network relevance of these features (Figure 5D and 5E). The majority of studies reported that increased time spent in strongly connected states was associated with preserved network integration and neural flexibility (often correlating with Healthy Controls), while a predominance of sparsely connected states characterized network disintegration and severe clinical phenotypes. Regarding state dynamics, a higher number of transitions was generally associated with preserved topological flexibility, thus better clinical outcomes, although two studies observed the inverse relationship.

**Figure 5.**
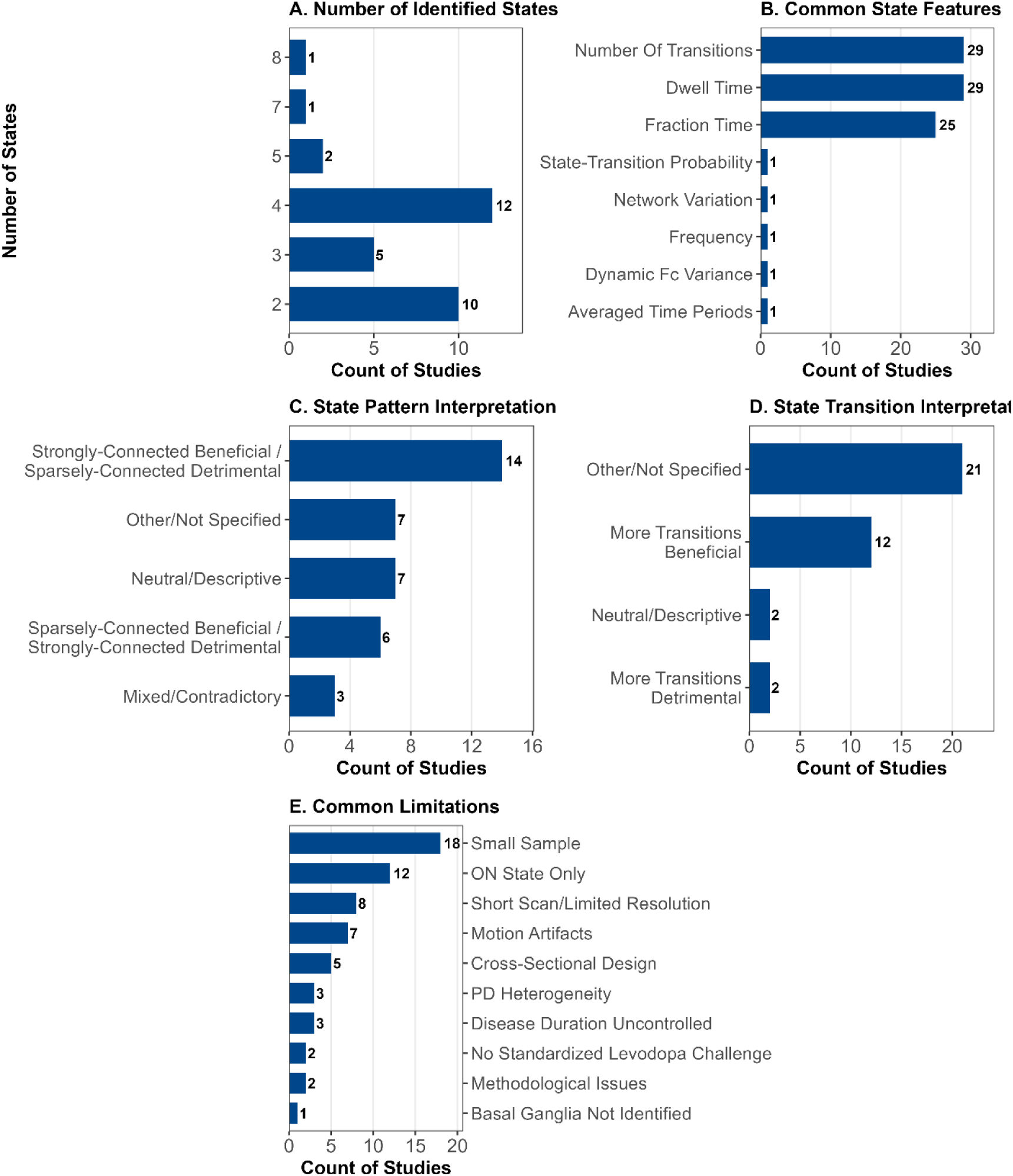
Variability in findings of the dFC analyses. (A) The number of dFC states identified across studies is variable, with two, three, and four states being the most common choices. (B) In contrast, there is a strong consensus on the dynamic features calculated, with dwell time, fraction time, and number of transitions being the most frequently reported metrics. (C) Clinical interpretations of state patterns are variable; while most studies concluded that strongly-connected states were beneficial, a smaller cohort found the counter argument that sparsely-connected states were beneficial. (D) Similarly, conclusions on state transitions show variability, though more transitions beneficial is the most frequent finding. (E) A clear consensus on limitations points to a shared path forward, with small sample size, head motion artifacts, and scanning in the ON State only being the most commonly cited challenges.

Finally, Figure 5E summarizes the limitations frequently cited across articles. Small sample size (see DynaPD *Study Overview* tab) and acquisition constraints – specifically short scan duration and low temporal resolution – were the most prevalent concerns. Furthermore, medication status was frequently noted as a confounding factor, as findings derived from patients in the ’ON’ state may not generalize to the unmedicated disease state. Conversely, studies examining unmedicated patients often cited head motion artifacts as a significant limitation. Additional challenges included study design constraints (e.g., cross-sectional nature) and the inherent clinical heterogeneity of the PD population. Readers are invited to explore how these study limitations are distributed across study design in the DynaPD

## 4. Discussion

In this scoping review, we systematically extracted a wide range of information, covering study metadata, experimental design, methodological decisions, and findings, of studies implementing dynamic FC analyses in PD samples. We then mapped this information in interactive visualization to enable the exploration of the research landscape and the variability of design, methodological decisions, and findings across studies. In this section, we discuss the outlook of the fields considering the research landscape presented in this review.

### 4.1. The need for longitudinal studies, longer scan, and bigger sample: data aggregation through federated learning

Our landscape analysis reveals a distinct prevalence of cross-sectional designs characterized by relatively small sample sizes and limited scan durations (Figure 3, Figure 4, and *Study Overview* tab in DynaPD for the sample sizes). While these studies provide valuable snapshots of disease states, they inherently fail to capture the temporal trajectory of PD progression. This limitation is particularly important in the context of precision neuroimaging, where the objective shifts from characterizing group-level differences to mapping individualized network deviations ^18,19^. Although dFC aims to quantify the temporal dynamics of brain activity, applying it to cross-sectional data restricts the analysis to the dynamics at a single time point. However, PD is a progressive neurodegenerative disorder where the network topology of individuals evolves over time. To elucidate the neural underpinnings of this evolution, study designs must capture not only the variability across individuals but also the dynamics within individuals across time ^20^.

The 10 studies in our review that did incorporate longitudinal or repeated-measures approaches offer empirical evidence for this necessity. The most direct support comes from the longitudinal studies of Cao et al. ^21^ and Li et al. ^22^, both of which show that systematic shifts in the dwell time and occupancy of specific dFC states correlated directly with worsening clinical trajectories, including autonomic, motor, and cognitive deterioration. Furthermore, repeated-measures designs have proven essential for capturing treatment sensitivity. Studies by Si et al. ^23^, Chen et al. ^24^, Liu et al. ^25^, Lee et al. ^26^, and Palmer et al. ^27^ reveal significant levodopa-associated within-subject alterations in state dwell times, transition probabilities, and network dynamics. Extending this to surgical interventions, Chu et al. ^28^ observed DBS-associated dFC remodeling, while Wu et al. ^29^ demonstrated that preoperative dynamic graph features could predict individual Subthalamic Nucleus-DBS responses. Notably, Boon et al. ^30^ added an important finding that dFC correlated with cognitive dysfunction cross-sectionally, but not longitudinally, illustrating that while powerful, the added value of repeated measures may be domain-specific. Taken together, these studies support the need for longitudinal and repeated-measures designs to separate state-dependent fluctuations from trait-level disease trajectories, a distinction that cross-sectional comparisons alone cannot achieve.

Beyond study design, the robustness of dFC estimates is heavily dependent on acquisition parameters. Previous methodological work suggests that reliable estimation of dynamic states requires sufficient scan duration, typically exceeding 10 to 15 minutes,^17^ although this should not be interpreted as a universal threshold independent of TR or analytical framework. Contrary to this recommendation, our review indicates that the majority of existing PD dFC studies employed scan lengths shorter than 10 minutes (Figure 4B; see also the Method Explorer tab in the DynaPD app). Furthermore, many studies utilized relatively long TR, resulting in low temporal resolution (Figure 4A). This combination of short scan duration and low temporal resolution compromises the fidelity of dFC analysis. First, insufficient sampling time increases the risk that identified states represent noise or transient artifacts rather than stable biological features ^31^. Second, dFC aims to capture neural reconfiguration occurring at the scale of seconds to milliseconds. High-TR acquisition acts as a temporal low-pass filter, potentially aliasing rapid neural fluctuations and obscuring the fine-grained transition patterns that may be most relevant to PD pathology. Although shorter TR can increase the temporal sampling density, it cannot compensate for short scan duration, as BOLD-fMRI remains constrained by the hemodynamic response. Moreover, simulation work on sliding-window correlation and k-means clustering has shown that state-transition and duration estimates are more strongly influenced by window length, window offset, noise, and filtering than by sampling rate alone ^32^. Therefore, future dFC studies in PD should avoid treating shorter TR and longer scan duration as interchangeable solutions. Instead, acquisition protocols should be optimized by jointly considering total acquisition time and temporal sampling density. Accelerated acquisition sequences, such as multiband imaging, may help improve sampling density, but their use in PD should take into account the potential reduction in subcortical signal-to-noise ratio associated with greater acceleration factors ^33^.

Moreover, our analysis of cohort demographics reveals an overrepresentation of males across the reviewed studies. The median proportion of male participants in the PD samples is around 60% (as shown in the *Study Overview* tab of the DynaPD app), which reflects the well-documented male prevalence of Parkinson’s disease^34^. Although these cohort proportions are representative of the clinical population, the specific neurobiological impact of sex on dynamic brain network reconfigurations is still a key area of research in this field. Given the well-documented clinical variations between men and women with regard to PD symptom presentation, progression rates and treatment responses, understanding how these phenotypic differences manifest in dynamic functional connectivity is essential. Future dFC research should consider sex as one of the primary biological variables to determine whether distinct topological trajectories exist between male and female patients.

Finally, the statistical power of the reviewed field is constrained by sample size, with most studies including fewer than 50 participants (see *Study Overview* in the DynaPD app). Recent work on brain-wide association studies has demonstrated that reproducible associations between neuroimaging phenotypes and behavioral traits require significantly larger datasets, often in the order of thousands ^35^. Achieving such numbers in clinical populations, such as PD, is challenging for individual sites due to recruitment and resource constraints. Our analysis shows that while individual sample sizes are small, the aggregate data spans multiple public and private sources (Figure 2C). This fragmentation presents an opportunity for data aggregation; however, pooling clinical data across sites is often hindered by privacy regulations and data governance issues. Federated Learning (FL) emerges as a promising solution to this challenge ^36^.

In an FL framework, model training occurs locally at each site (e.g., identifying dFC states on local private data), and only the model updates (gradients or parameters) are aggregated centrally. This approach would allow the PD research community to leverage the statistical power of large-scale, multi-site data to construct robust dFC biomarkers while strictly adhering to patient privacy and data protection standards.

### 4.2. Advancement in methods with the consideration of clinical interpretation

Our analysis found an important methodological convergence within the field: the majority of studies employ a pipeline combining sliding-window correlation with k-means clustering (Figure 5E). It is important to note that this methodological convergence is not without controversy. The dFC field more broadly faces unresolved debates surrounding the statistical validity of sliding-window estimates, the sensitivity of derived states to arbitrary parameter choices and their physiological interpretability^37^. While this standardization facilitates comparisons across cohorts, it relies on the rigid selection of a fixed window length, which constitutes a significant researcher degree of freedom. This imposition of a static temporal scale risks obscuring transient neural events that do not align with the chosen window, potentially oversimplifying the true underlying dynamics. Alternative data-driven approaches, such as Hidden Markov Models (HMM) ^38^ or Co-Activation Patterns (CAPs) ^39^, avoid these arbitrary windowing parameters and may offer a more flexible representation of neural state transitions. However, the adoption of these mathematically complex models must be carefully balanced against biological plausibility and clinical interpretability. For dFC markers to translate into clinical practice, they must remain grounded in concepts that clinicians can interpret rather than abstract statistical probabilities.

Once states are identified, the subsequent characterization of PD pathology relies on temporal features related to the states, specifically number of transitions, dwell time, and fraction time (Figure 5B). While these metrics robustly capture the temporal stability of brain states, they offer limited insight into the topological organization within those states. Given that PD is fundamentally a network disorder where motor and cognitive deterioration are intertwined, relying solely on temporal metrics may be insufficient ^7^. A subset of studies in our review incorporated graph theoretical measures (e.g., global or local efficiency; see *Findings Explorer* in DynaPD). This approach is arguably more biologically plausible for elucidating the mechanisms of PD, as it directly quantifies the efficiency of information transfer between distributed systems ^4^. Future work should prioritize these topological features to better map, for example, the compensatory hypothesis of PD cognitive decline, where cognitive decline results from compensating for deterioration in motor-related brain networks. However, as also stated above the use of these higher order metrics must remain balanced with the need for clear clinical interpretation.

Finally, regarding the goal of precision neuroimaging, the dominant analytical strategy involves concatenating data to identify common states across the group, which are then back-projected to individuals. While effective for identifying general disease phenotypes, this group-to-individual approach risks masking subject-specific state configurations that may be important for personalized diagnosis ^19^. In the context of precision medicine, a bottom-up approach is preferable: first identifying stable states at the individual level, and subsequently examining their commonality across the population. This strategy allows for the preservation of unique individual trajectories – an important factor for tailoring patient-specific treatments (e.g., DBS targeting). However, as noted in Section 4.1, the successful implementation of this individualized approach is contingent upon the acquisition of high-density, long-duration scans for each patient.

### 4.3. Neural Flexibility and Clinical Phenotype: Linking Brain Dynamics to Disease Severity

Our findings also reveal a consistent dynamic phenotype in PD: a temporal shift favoring sparsely connected (segregated) states over densely connected (integrated) ones, accompanied by a reduction in state transitions relative to HC (Figure 5C and 5D). These findings align with previous study^40^, suggesting that the dopamine loss in PD is mirrored by a loss of neural flexibility.

From a network perspective, strongly and densely connected states are metabolically costly but essential for global information integration. High-level cognitive tasks and complex motor planning require the brain to transiently shift from a segregated architecture into a globally integrated state to facilitate widespread information transfer ^41^. The inability of the PD brain to access or sustain these strongly connected states and to perform transition between states likely reflect a failure in this global integration mechanism. Without the ability to transiently integrate distributed brain regions, patients struggle with tasks requiring coordination between sensorimotor and associative networks, manifesting clinically as deficits in executive function and complex motor sequencing ^40^.

Clinically, the studies included in this review collectively point to a potential link between dFC alterations and symptom severity in PD. Across several cohorts, greater time spent in sparsely connected states (or reduced dwell time in densely and strongly connected states) and fewer transitions were associated with worse clinical profiles, including higher MDS-UPDRS III scores^42^, the presence of freezing of gait ^25^, and greater tremor severity^43^. Although less frequently examined, several studies also reported similar associations for non-motor symptoms, such as rapid eye movement sleep behavior disorder^44^, and cognitive impairment^10,45^.

The mechanistic basis of these associations may be partly attributed to dopaminergic disruption as a central contributor to dFC alterations in PD. A previous study demonstrated that dFC features correlate with striatal dopaminergic loss, as measured by dopamine transporter scan, suggesting a link between impaired large-scale functional dynamics and dopamine depletion ^42^. Complementing this, studies investigating medication effects indicate that dopamine replacement therapy tends to increase neural flexibility, reflected by more frequent state transitions ^26^. Extending this to the topological level, graph-theoretical analyses have further demonstrated that levodopa normalizes the organization of the disrupted rich-club network in PD. This suggests that dopaminergic modulation acts on both state transition dynamics and the hierarchical architecture of large-scale functional brain networks ^46^. Together, these findings suggest that dFC metrics may capture clinically meaningful aspects of PD pathophysiology.

Despite these converging observations, greater dwell time in sparsely connected states and lower transitions cannot be generally concluded as a common clinical feature of PD, given the notable discrepancies across studies. For example, a study reported the opposite pattern, showing that lower dwell time in sparsely connected states was associated with higher motor severity ^47^. Similarly, another work found increased state transitions in PD patients with dementia^48^. Even within individual studies, results may diverge: the association between dopaminergic integrity and dFC features was present in only one of the two cohorts they examined^42^. These inconsistencies underscore that dFC-clinical relationships are not yet stable across methodological choices, disease stages, or patient characteristics, highlighting the need for greater harmonization and longitudinal designs.

### 4.4. Report standardization

A substantial challenge encountered during the systematic extraction of data for this review was the inconsistency in reporting practices across studies. This variability was most pronounced in the description of the MRI data preprocessing pipelines and sample characteristics (e.g., incomplete reporting of medication state, disease severity, disease duration, or specific motor subtype). Such reporting heterogeneity introduces researcher degrees of freedom that are often undisclosed, making direct replication challenging and hindering the precise assessment of methodological validity. Furthermore, inconsistencies in the nomenclature used to label large-scale brain networks were observed across the reviewed studies. This represents an additional source of methodological variability that complicates synthesizing and comparing findings across studies. Similar challenges have been identified in the broader network neuroscience literature, underscoring the need for more consistent network definitions and reporting practices ^49^.

To mitigate this, it is recommended to move beyond ad-hoc descriptions and adopt rigorous and standardized reporting frameworks. For neuroimaging parameters, adherence to guidelines such as those proposed by the Committee on Best Practice in Data Analysis and Sharing (COBIDAS) is recommended ^50^. Specifically, studies should utilize standardized checklists to report acquisition sequences, preprocessing software versions, and critical denoising steps. Furthermore, regarding clinical phenotyping, we suggest a consensus on reporting MDS-UPDRS sub-scores rather than aggregate totals alone, as well as the explicit documentation of Levodopa Equivalent Daily Dose (LEDD) and medication state (ON/OFF) during scanning ^51^. Without this granular clinical data, it is challenging to disentangle whether dFC alterations are driven by disease pathology or pharmacological effects.

Moreover, report standardization is a prerequisite for the future of automated knowledge synthesis. As the volume of literature grows, manual extraction becomes increasingly difficult. Standardized, machine-readable reporting formats (e.g., structured abstracts or Brain Imaging Data Structure-compliant metadata) would facilitate the use of Natural Language Processing (NLP) and Artificial Intelligence (AI)-driven tools to automatically extract methodological details for living systematic reviews ^52,53^. Importantly, recent perspective emphasize that to overcome translational bottlenecks, the neuroimaging research must evolve beyond static data repositories, by establishing a dynamic, empirically guided metascientific framework to actively steer method development ^54^. Furthermore, as discussed in Section 4.1, the implementation of FL relies heavily on data harmonization. Standardizing the input features (clinical and imaging data) is the first step toward creating the decentralized and interoperable datasets necessary to train robust, privacy-preserving AI models for PD.

### 4.5. Limitations

The present study is not without limitations. First, our review was restricted to fMRI-derived dFC, excluding studies utilizing other neuroimaging modalities such as Electroencephalography (EEG) or Magnetoencephalography (MEG). While fMRI offers superior spatial resolution, electrophysiological methods provide direct measures of neural activity with millisecond precision. Given that dFC phenomena may manifest across multiple spatiotemporal scales, future integrative reviews synthesizing findings across these modalities are necessary to achieve a complete understanding of neural dynamics in PD. Second, while we acknowledge the clinical heterogeneity of PD, our synthesis largely treated PD as a general category. Although we identified studies focusing on specific subtypes (e.g., tremor-dominant vs. postural instability-gait disorder), the aggregate nature of a landscape review may obscure distinct dynamic signatures unique to these subgroups. As the field matures and more subtype-specific data becomes available, more granular meta-analyses will be required to disentangle general disease effects from phenotype-specific network alterations. Finally, the DynaPD application is currently founded exclusively on the dataset extracted for this review. While this ensures high data quality through manual curation, it limits the immediate scope of the tool. However, the functionality of the app can be readily expanded, and the underlying database can be continuously updated – potentially through automated NLP pipelines as discussed in Section 4.4 – to incorporate emerging studies and additional analytical dimensions not currently considered. By open-sourcing this tool, we invite the community to contribute to this living evidence synthesis, ensuring it remains a relevant resource for the field.

### 4.6. Conclusion and outlook

This systematic review elucidates the rapidly evolving yet fragmented landscape of dynamic FC research in PD. Our synthesis reveals that while the field has converged on a core set of analytical techniques, it remains characterized by substantial heterogeneity in preprocessing strategies and variability in research findings. Consequently, future studies must prioritize experimental designs that overcome previous constraints – specifically by adopting longitudinal sampling and extended acquisition protocols – while adhering to standardized reporting guidelines to mitigate analytical degrees of freedom.

Beyond the static evidence synthesized herein, the accompanying DynaPD application serves as a dynamic, living resource for the community. By enabling interactive exploration of the data, the tool allows researchers to navigate the complex methodological landscape and contextualize their own findings within the broader field. Looking forward, the application is developed for scalability; as the community moves toward standardized reporting, the platform can be extended to leverage automated information extraction. This integration of systematic review and interactive software aims to transform a diverse body of literature into a coherent, cumulative science, accelerating the translation of dFC biomarkers into precision neurology.

## Data Availability

The dataset generated during the current study, which includes the comprehensive information extracted from the 37 eligible articles, is available in the GitHub repository [https://github.com/kristantodan12/dFC-PD] under the filename Database.xlsx. All data underlying the figures and the interactive visualizations in the DynaPD application are contained within this file.

## Code Availability

The source code developed for the DynaPD interactive application and the associated data visualization scripts are available in the GitHub repository [https://github.com/kristantodan12/dFC-PD]. The web-based version of the application is also accessible for immediate use. The link is available in the GitHub repository.

## Acknowledgment

DK was supported by the Young Researchers’ Fellowship from the University of Oldenburg and by a grant from the German Research Foundation (DFG) to Andrea Hildebrandt (HI 1780/7-1) and Carsten Gießing (GI 682/5-1) as part of the DFG priority program “META-REP: A Metascientific Program to Analyze and Optimize Replicability in the Behavioral, Social, and Cognitive Sciences” (SPP 2317). AA was funded by DFG as part of the Research Training Group (RTG)-2783 (Project ID: 456732630).

## CRediT authorship contribution statement

DK: conceptualization, data curation, methodology, software, formal analysis, visualization, writing – original draft, writing – review and editing. DC: data curation, formal analysis, writing – review and editing. AA: conceptualization, formal analysis, writing – review and editing.

